# Sex-specific Associations Between Traumatic Experiences and Resting-state Functional Connectivity in the Philadelphia Neurodevelopmental Cohort

**DOI:** 10.1101/2020.12.18.423455

**Authors:** Shiying Wang, Jeffrey G. Malins, Heping Zhang, Jeffrey R. Gruen

**Affiliations:** Department of Biostatistics, Yale University School of Public Health, New Haven, CT, USA; Department of Psychology, Georgia State University, Atlanta, GA, USA; Haskins Laboratories, New Haven, CT, USA; Departments of Pediatrics and Genetics, Yale University School of Medicine, New Haven, CT, USA

**Keywords:** Functional connectivity, traumatic experiences, sex differences, somatomotor network, default mode network

## Abstract

**Background:** Traumatic experiences during childhood or adolescence are a significant risk factor for multiple psychiatric disorders and adversely affect cognitive functions. Resting-state functional magnetic resonance imaging has been used to investigate the effects of traumatic experiences on functional connectivity, but the impact of sex differences has not been well documented. This study investigated sex-specific associations between resting-state functional connectivity and traumatic experiences in typically developing youth.

**Methods:** The sample comprised 1395 participants, ages 8 to 21 years, from the Philadelphia Neurodevelopmental Cohort. Resting-state functional connectivity was characterized by voxel-wise intrinsic connectivity distribution parameter values derived from resting-state functional magnetic resonance imaging. Traumatic experiences were assessed based on a structured psychiatric evaluation. Sex, the number of traumatic events, and their interaction were regressed onto voxel-wise intrinsic connectivity distribution parameter values. Brain regions that passed cluster correction were used as seeds to define resting-state networks.

**Results:** After quality control, the final sample included 914 participants (mean (SD) age, 14.6 (3.3) years; 529 (57.8%) females; 437 (47.8%) experienced at least one kind of traumatic event). Four discrete anatomical clusters showed decreased functional connectivity as the number of traumatic events increased. The resting-state networks defined by using these four clusters as seeds corresponded with the somatomotor network. Sex-specific associations were identified in another four clusters for which males showed increased connectivity, and females showed decreased connectivity as the number of traumatic events increased. The resting-state networks defined by the four sex-specific clusters corresponded with the default mode network.

**Conclusions:** Traumatic experiences are associated with an alteration of resting-state functional connectivity in the somatomotor network in youth without psychiatric diagnoses. The associations differ in direction between males and females in the default mode network, suggesting sex-specific responses to early exposure to trauma.

## Introduction

Traumatic experiences during childhood or adolescence, defined as exposure to actual or threatened death, serious injury, or sexual abuse (Association, 2013), are a significant risk factor for multiple psychiatric disorders, such as post-traumatic stress disorder (PTSD), major depressive disorder (MDD), and substance abuse (Copeland, Keeler, Angold, & Costello, 2007; Heim & Nemeroff, 2001; Kilpatrick et al., 2003). Traumatic experiences are associated with adverse effects on cognitive function including processing speed, memory and language skills, as well as childhood and adolescent development (Majer, Nater, Lin, Capuron, & Reeves, 2010; Sylvestre, Bussières, & Bouchard, 2016). In the U.S., the prevalence of traumatic experiences during childhood and adolescence ranges from 10.14% to 23.3% (Briere & Elliott, 2003; Pérez-Fuentes et al., 2013). However, the biological mechanisms underlying the associations between traumatic experiences and psychopathology are not well understood.

Functional magnetic resonance image (fMRI) is a powerful tool for analyzing the neurobiology of cognitive functioning. In particular, resting-state fMRI can delineate intrinsic functional brain networks enabling auditory and visual processing, attention, memory retrieval, and many of the critical functions collectively called cognition. Resting-state functional connectivity (rsFC), which examines inter-correlations of activity between brain regions over time, has been utilized to investigate the impact of trauma on brain function. Compared to controls, PTSD is associated with decreased rsFC within the default mode network (DMN; Bluhm et al., 2009; Lanius et al., 2010; Sripada et al., 2012; Viard et al., 2019), increased rsFC within the salience network (SN), and elevated cross-network connectivity between both networks (Sripada et al., 2012). Traumatic exposure in non-PTSD is associated with alterations within the DMN (Lu et al., 2017) and insula-based networks (Marusak, Etkin, & Thomason, 2015). These findings suggest that traumatic experiences play a central role in rsFC changes in both PTSD and non-PTSD individuals. However, the sample sizes of these association studies are small and lack power to stratify for sex and other covariates.

Female victims are more likely than males to develop PTSD after experiencing trauma (Breslau, 2009; Tolin & Foa, 2008), and there are sex differences in neural activation. With an fMRI fear-perception paradigm, when compared to males, females with PTSD show greater activity in the dorsal brainstem and less activity in the hippocampus (Felmingham et al., 2010). In a fear learning and extinction paradigm, males with PTSD increase activation compared to females, in the left rostral dorsal anterior cingulate cortex during extinction recall (Shvil et al., 2014). Based on these and other studies, we hypothesize that there are sex-specific associations between traumatic exposure and rsFC, which can partially account for the observed differences in rates of PTSD and qualitative differences in clinical presentation between males and females (Felmingham et al., 2010; Shvil et al., 2014).

## Methods

### Participants

We tested our hypotheses in the Philadelphia Neurodevelopmental Cohort (PNC; Satterthwaite et al., 2016), a large non-psychiatric sample of youth ages 8 to 21 years. Data were collected through a collaborative study between the Center for Applied Genomics at Children’s Hospital of Philadelphia and the Brain Behavior Laboratory at the University of Pennsylvania from 2009 to 2011. For this study, we analyzed a subsample of 1395 participants with both structured psychiatric evaluations and resting-state fMRI brain scans.

### Ethical Considerations

Appropriate informed consent was obtained from all study participants. The University of Pennsylvania and the Children’s Hospital of Philadelphia approved all study procedures (Satterthwaite et al., 2016).

### Traumatic experience assessment

Each participant received a structured screening interview, called GOASSESS (Calkins et al., 2014), based on a modified version of the Kiddie-Schedule for Affective Disorders and Schizophrenia (Kaufman et al., 1997). We used nine questions from the PTSD assessment scale (TableS1) to assess traumatic experiences (Barzilay et al., 2019). Responses for each question were assigned a value of 1 for yes, and 0 for no (Appendix S1). For each participant, we summed the number of unweighted traumatic events (TE) for use in subsequent regression analyses.

### Image acquisition and preprocessing of MRI data

MRI images were collected using a single 3T Siemens TIM Trio whole-body scanner located in the Hospital of the University of Pennsylvania (Satterthwaite et al., 2014). The acquisition of anatomical images and resting-state fMRI are detailed in Appendix S2. We preprocessed resting-state fMRI data in AFNI (Cox, 1996). The first four TRs were removed, followed by slice-scan time correction, alignment with anatomic images, and registration to MNI152 space. We corrected for head motion by aligning to the volume with the minimum outlier fraction, smoothed images using a 6 mm FWHM Gaussian kernel, and performed a general linear model using six motion parameters and their derivatives as regressors as well as frequency components

between 0.01-0.08 Hz. Volumes were censored if they exceeded 0.3 mm point-to-point Euclidean movement or had >10% outlier voxels. Then, we performed tissue-based regression to regress out the average signals of individual eroded white matter masks and the first three principal components of individual eroded lateral ventricle masks detailed in Appendix S3.

### Quality control

After preprocessing of fMRI data, we removed samples with average motion per TR >0.3 mm or censor fractions >30% (n=225). Samples whose functional and anatomic images did not co-register properly (based on visual inspection) were removed (n=69). Based on self-reported medical history, samples with serious brain injury were excluded (n=178). In addition, two samples were excluded for missing traumatic experience records, and seven were excluded for missing data concerning maternal education. Data from 914 participants passed quality control procedures and were used in the primary analyses.

### Intrinsic connectivity distribution (ICD)

We used ICD to measure voxel-level functional connectivity as previously described (Scheinost et al., 2012). ICD is a data-driven method that does not require prior information to define regions of interest or an arbitrarily chosen correlation threshold. For each voxel, a histogram of correlation coefficients was generated to characterize the rsFC between that voxel and all other voxels. These histograms were approximated using a function with two parameters, alpha and beta (Appendix S4). Smaller alpha values and larger beta values for any voxel represent a higher density of strong connections with other voxels. We computed ICD parameter values for voxels within an MNI152 gray matter mask (31,053 voxels) for each participant using custom scripts in R (version 3.6.0).

### Regression on ICD parameter values

We conducted regression analyses on subject-wise ICD parameter values to define seeds for functional connectivity analysis. ICD parameter values were smoothed using a 4 mm FWHM Gaussian kernel. In the regression model for each voxel, ICD parameter values were regressed on the number of TE’s that participants experienced. Covariates included sex, age, and the number of years of maternal education. To test for sex-specific associations, interaction terms between sex and TE number were also included in the model. ICD alpha and beta values were respectively analyzed. Cluster correction was performed with the voxel-wise threshold set at *p* = 0.001, cluster corrected at *α*= 0.05.

### Seed-based functional connectivity analysis

We computed correlation maps using the clusters that passed correction as seed regions. Correlation coefficients between the time series of a seed region and each voxel were calculated and transformed to z scores. Then, using a *t*-test, we tested whether z scores were significantly different from 0. Regions passing the thresholds of *p* < 1×10^−44^ and FDR < 3×10^−16^ were defined as a resting-state network. The correlation maps were drawn using SUMA (Saad, Reynolds, Argall, Japee, & Cox, 2004). Identified networks were compared with seven well-defined reference networks (Thomas Yeo et al., 2011) to determine the percentage of voxels overlapping with each reference network. The parcellation of reference networks was downloaded from https://surfer.nmr.mgh.harvard.edu/fswiki/CorticalParcellation_Yeo2011.

## Results

### Demographics

Of 914 participants with mean (SD) age 14.6 (3.3) years, 529 (57.8%) were females, 396 (43.3%) were white, and 437 (47.8%) experienced at least one kind of TE. Table 1 shows the demographic information for participants by number of TE’s. Participants who experienced more TE’s were older and had fewer years of maternal education. The demographic information of participants with and without traumatic exposure, as well as the frequencies of participants in each category of TE, are provided in TableS2 and TableS3.

**Table 1.** Demographic information for participants by number of traumatic events

|  | Participants (N=914) |  |  |  |  |  |  |
| --- | --- | --- | --- | --- | --- | --- | --- |
| Characteristics | TE=0<br>(n=477) | TE=1<br>(n=219) | TE=2<br>(n=113) | TE=3<br>(n=58) | TE=4<br>(n=36) | TE=5<br>(n=7) | TE=6<br>(n=4) |
| Female sex, No.<br>(%) | 274<br>(57.4) | 133<br>(60.7) | 68 (60.1) | 31 (53.4) | 17 (47.2) | 3 (42.8) | 3<br>(75.0) |
| Race |  |  |  |  |  |  |  |
| African-American, No.<br>(%) | 187<br>(39.2) | 99 (45.2) | 66 (58.4) | 39 (67.2) | 24 (66.7) | 5 (71.4) | 3<br>(75.0) |
| European-American, No.<br>(%) | 233<br>(48.8) | 97 (44.3) | 45 (39.8) | 12 (20.7) | 7 (19.4) | 1 (14.3) | 1<br>(25.0) |
| Other, No. (%) | 54 (11.3) | 22 (10.0) | 2 (1.8) | 7 (12.1) | 5 (13.9) | 1 (14.3) | 0 (0) |
| Age, mean<br>(SD), y | 13.9<br>(3.5) | 14.9<br>(3.0) | 15.2<br>(2.8) | 15.7<br>(3.1) | 16.6<br>(2.6) | 16.6<br>(3.1) | 17.8<br>(1.7) |
| ME, mean<br>(SD), y | 14.4<br>(2.4) | 14.0<br>(2.5) | 14.1<br>(2.6) | 13.3<br>(2.0) | 13.7<br>(2.3) | 14.7<br>(1.7) | 12 (0) |

Abbreviations: TE, traumatic events; ME, years of maternal education

### Regression on ICD parameter values

Using resting-state fMRI data, we computed ICD parameter values for voxels within a gray matter mask for each participant. Then, we conducted a regression analysis on ICD parameter values to analyze the main effect of TE number and the interaction between sex and TE number. Results reported here are for the alpha parameter.

After cluster correction, we identified four clusters that showed a significant main effect of TE number on ICD alpha values (Table 2, Clusters 1-4). Within these clusters, as TE number increased, alpha values increased (Figure 1(a)), which means that among participants with higher TE exposure, voxels in these regions had reduced functional connectivity with other voxels.

**Table 2.**
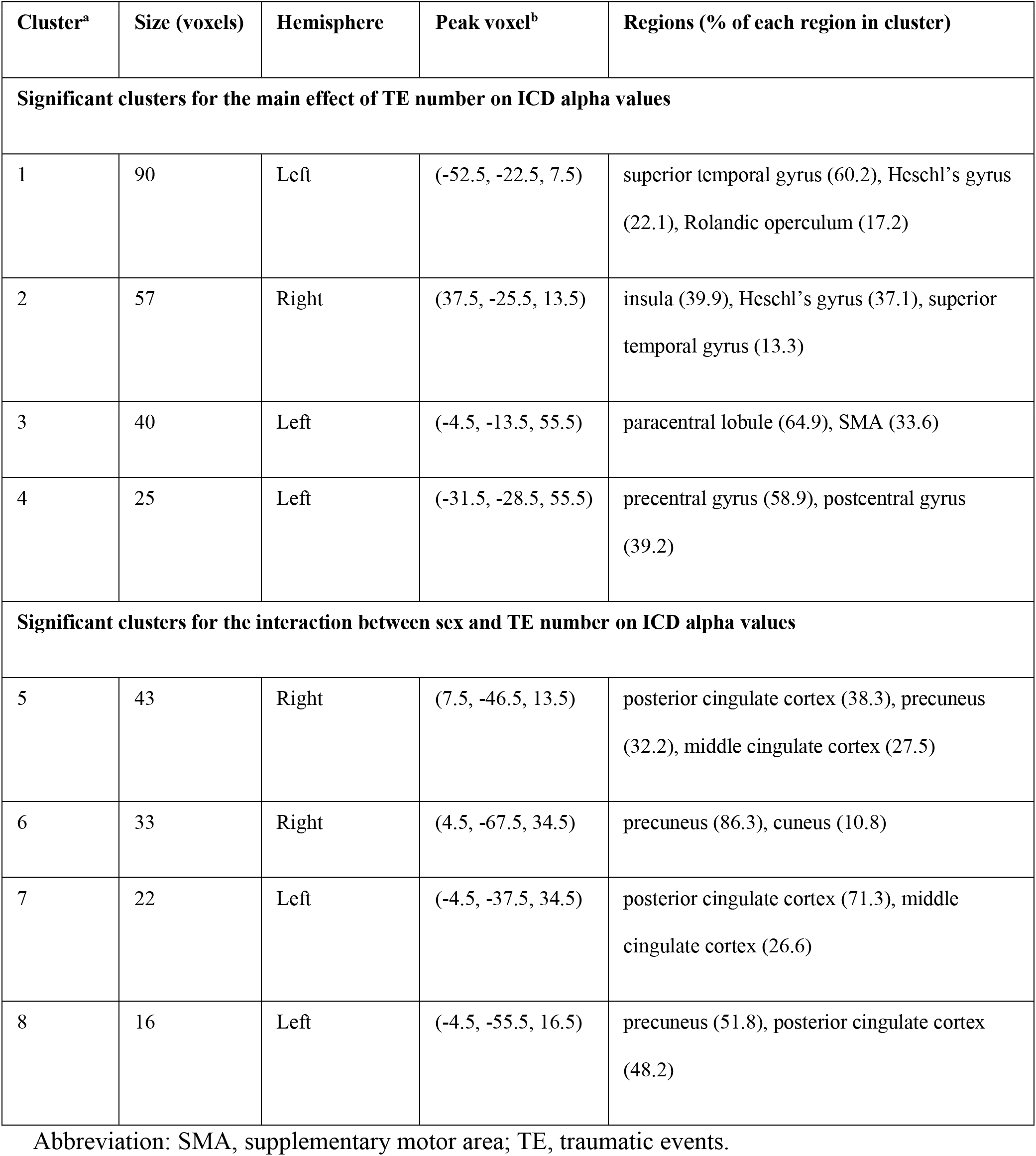
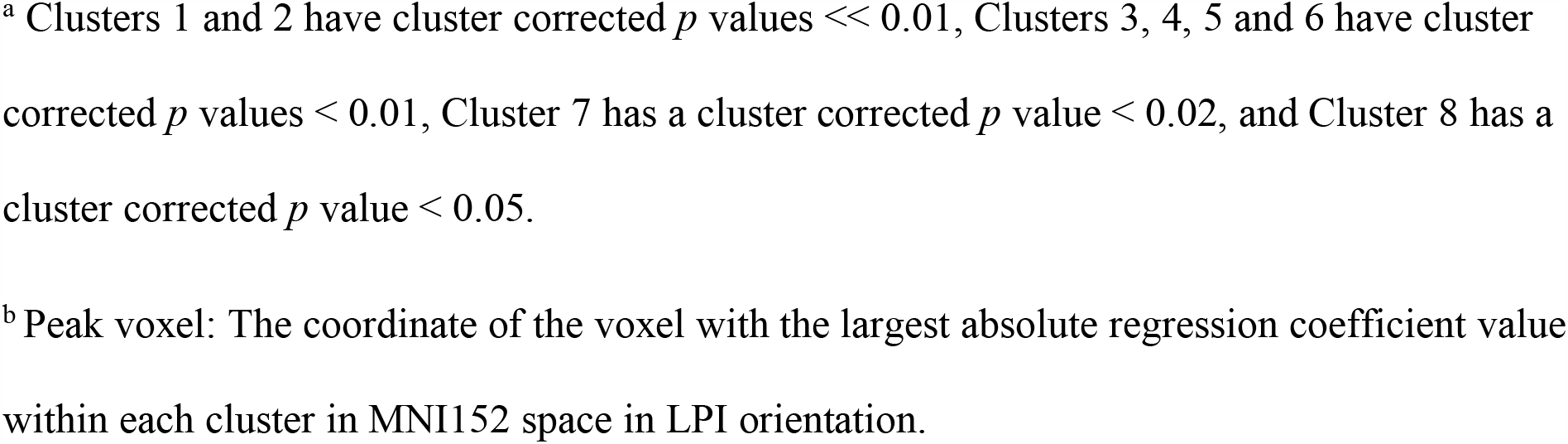
Summary of significant clusters that passed cluster correction in regression on ICD alpha values

| Cluster<sup>a</sup> | Size (voxels) | Hemisphere | Peak voxel<sup>b</sup> | Regions (% of each region in cluster) |
| --- | --- | --- | --- | --- |
| **Significant clusters for the main effect of TE number on ICD alpha values** |  |  |  |  |
| 1 | 90 | Left | (-52.5, -22.5, 7.5) | superior temporal gyrus (60.2), Heschl's gyrus (22.1), Rolandic operculum (17.2) |
| 2 | 57 | Right | (37.5, -25.5, 13.5) | insula (39.9), Heschl's gyrus (37.1), superior temporal gyrus (13.3) |
| 3 | 40 | Left | (-4.5, -13.5, 55.5) | paracentral lobule (64.9), SMA (33.6) |
| 4 | 25 | Left | (-31.5, -28.5, 55.5) | precentral gyrus (58.9), postcentral gyrus (39.2) |
| **Significant clusters for the interaction between sex and TE number on ICD alpha values** |  |  |  |  |
| 5 | 43 | Right | (7.5, -46.5, 13.5) | posterior cingulate cortex (38.3), precuneus (32.2), middle cingulate cortex (27.5) |
| 6 | 33 | Right | (4.5, -67.5, 34.5) | precuneus (86.3), cuneus (10.8) |
| 7 | 22 | Left | (-4.5, -37.5, 34.5) | posterior cingulate cortex (71.3), middle cingulate cortex (26.6) |
| 8 | 16 | Left | (-4.5, -55.5, 16.5) | precuneus (51.8), posterior cingulate cortex (48.2) |

Abbreviation: SMA, supplementary motor area; TE, traumatic events.
<sup>a</sup> Clusters 1 and 2 have cluster corrected $p$ values $\ll 0.01$ , Clusters 3, 4, 5 and 6 have cluster corrected $p$ values $< 0.01$ , Cluster 7 has a cluster corrected $p$ value $< 0.02$ , and Cluster 8 has a cluster corrected $p$ value $< 0.05$ .
<sup>b</sup> Peak voxel: The coordinate of the voxel with the largest absolute regression coefficient value within each cluster in MNI152 space in LPI orientation.

**Figure 1.**
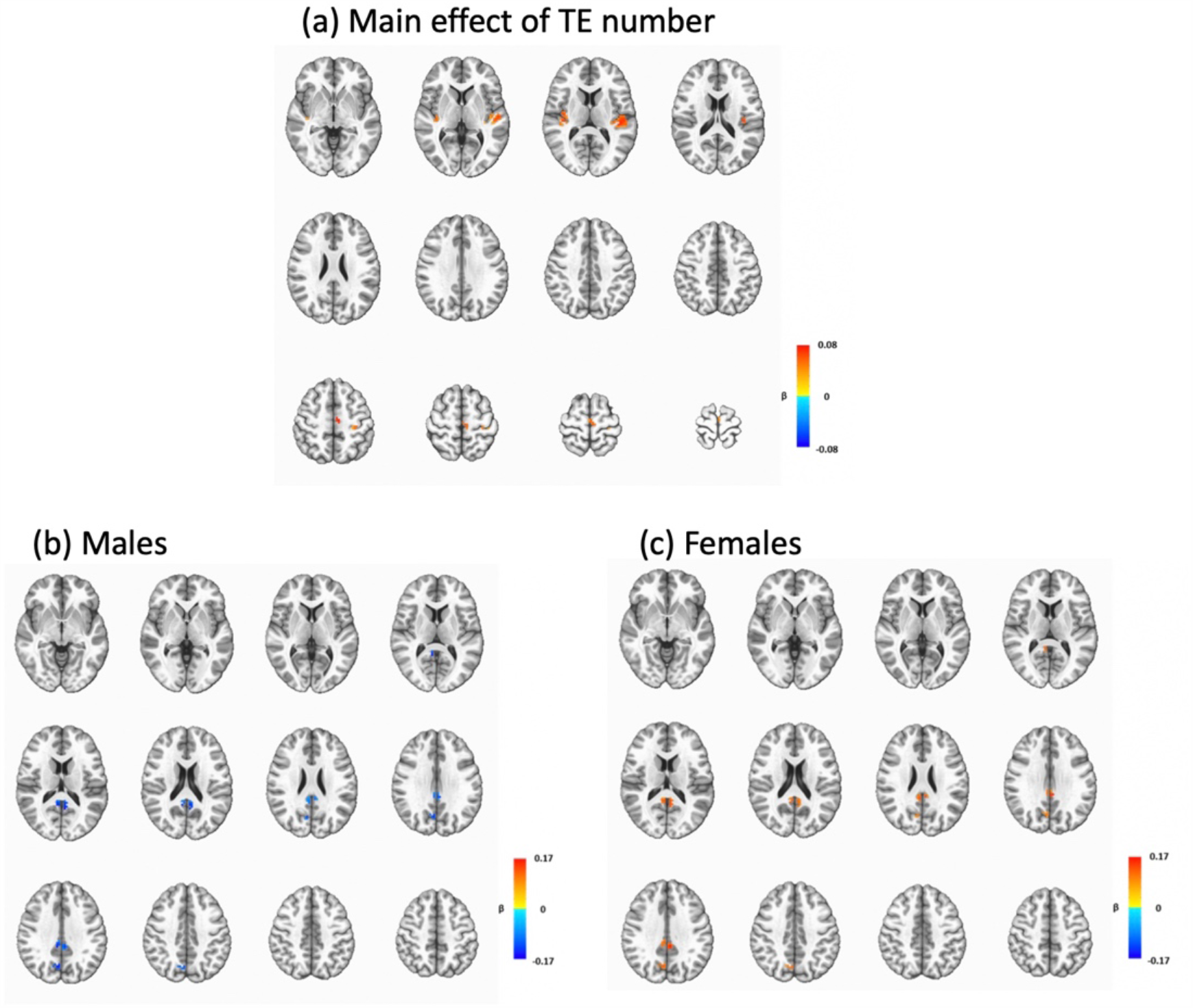
Significant clusters that passed cluster correction in regression on ICD alpha values. (a) Significant clusters for the main effect of TE number on ICD alpha values. Four clusters passed cluster correction (voxelwise *p* < 0.001, cluster corrected at *p* < 0.05). Colors represent the value of the regression coefficient of TE number for voxels within significant clusters. The space between layers is 7 mm. (b) (c) Significant clusters for the interaction between sex and the number of traumatic events. Four clusters passed cluster correction (voxelwise *p* < 0.001, cluster corrected at *p* < 0.05). To better visualize divergent effect directions for females and males in these clusters, the regression coefficients of the main effect term (TE number) and the interaction term (sex*TE number) in the regression model were integrated. Colors represent the amount of ICD alpha value change as TE number increased one unit for males (b) and females (c). The space between layers is 5 mm.

We identified another four clusters that showed a significant interaction between sex and TE number on ICD alpha values (Table 2, Clusters 5-8). Within these clusters, males and females had divergent changes in rsFC as TE number increased. For males, the ICD alpha value decreased (Figure 1(b)), whereas for females, the ICD alpha value increased as TE number increased (Figure 1(c)). Within these specific clusters, males showed increased connectivity, whereas females showed decreased connectivity as TE number increased.

### Seed-based functional connectivity analysis

To identify correspondences between clusters and resting-state networks, we used all eight clusters as seed regions to construct correlation maps and to define eight resting-state networks.

Using Cluster 1 as a seed region, we constructed resting-state Network 1. It contained the bilateral superior temporal gyrus, Heschl’s gyrus, supplementary motor area (SMA), middle cingulate cortex, precentral gyrus, postcentral gyrus, Rolandic operculum and insula, and the left paracentral lobule (Figure 2(a), FigS1). We compared Network 1 with seven well-defined resting-state networks (Thomas Yeo et al., 2011). Among the total of 1922 voxels in Network 1, 82.6% overlapped with the somatomotor network (Table 3), which included somatomotor and auditory cortices. Network 1 included most of the cortical regions in Networks 2-4. Results for Networks 2-4 are presented in FigS2, FigS3, FigS4, TableS4, TableS5, and TableS6.

**Table 3:** Percentage of Network 1 and Network 5 that overlapped with the reference networks in Yeo et al. (2011).

| Network | Number of overlapping voxels in<br>Network 1 (%) | Number of overlapping voxels in<br>Network 5 (%) |
| --- | --- | --- |
| Visual | 0 (0) | 280 (16.1) |
| Somatomotor | 1588 (82.6) | 7 (0.4) |
| Dorsal attention | 1 (0.05) | 36 (2.1) |
| Ventral attention | 605 (31.5) | 67 (3.9) |
| Limbic | 0 (0) | 3 (0.1) |
| Frontoparietal | 0 (0) | 419 (24.1) |
| Default mode | 2 (0.1) | 1404 (80.9) |

**Figure 2.**
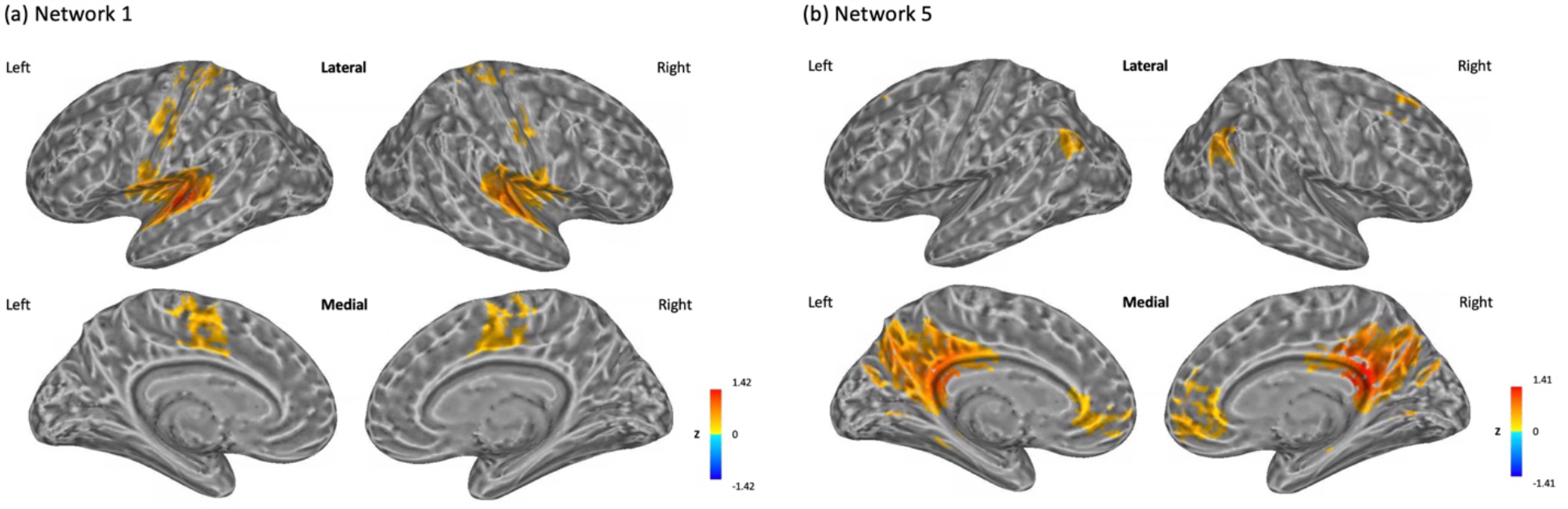
Resting-state Network 1 and resting-state Network 5. (a) Resting-state Network 1 defined by using Cluster 1 as a seed region. (b) Resting-state Network 5 defined by using Cluster 5 as a seed region. The correlation coefficients between the average time-series data of the seed region with every other voxel were calculated and transformed to z scores. A *t*-test identified significant regions (p < 1*10^−44^, FDR < 3*10^−16^) that defined a resting-state network. Colors represent the value of z scores within defined resting-state networks.

Using Cluster 5 as a seed region, we constructed resting-state Network 5. It contained the bilateral medial-orbital gyrus, anterior cingulate cortex, angular gyrus, precuneus, middle cingulate cortex, posterior cingulate cortex, middle frontal gyrus and cuneus, the left middle occipital gyrus, and the right superior frontal gyrus (Figure 2(b), FigS5). We compared Network 5 with reference resting-state networks (Thomas Yeo et al., 2011). Among 1735 voxels in Network 5, 80.9% overlapped with the DMN (Table 3). Network 5 included most of the cortical regions in Networks 6-8. Results for Networks 6-8 are presented in FigS6, FigS7, FigS8, TableS7, TableS8, and TableS9.

## Discussion

In a large cohort of non-psychiatric youth, we investigated sex-specific associations between traumatic experiences and rsFC. We identified four anatomical clusters that showed decreased rsFC with other brain regions as the number of TE’s increased. The defined resting-state networks using these clusters as seeds corresponded with the somatomotor network. We identified another four brain clusters that showed an effect of TE number, but in opposite directions for males and females. The defined resting-state networks using these clusters as seeds highly corresponded with the DMN.

### Main effect of TE number

The brain regions that showed significant associations between traumatic experiences and rsFC are similar to those identified in previous studies of adults with traumatic exposure (Lu et al., 2017; Philip, Kuras, et al., 2013) and patients with trauma-related psychiatric disorders, such as PTSD and MDD (De Bellis et al., 2002; Liu et al., 2010; Lorenzetti, Allen, Fornito, & Yücel, 2009; Sui et al., 2010; Tursich et al., 2015; Zhang et al., 2016).

Clusters 1 and 2 were mainly centered around temporal gyri, including the posterior temporal gyrus and Heschl’s gyrus. Temporal gyri play an essential role in auditory processing for receptive and expressive language. Wernicke’s area, located in the posterior portion of the left superior temporal gyrus, is critical for speech processing and language comprehension. In a small study of 48 young adults (average age 21.8 years), decreased regional homogeneity was observed in the bilateral posterior temporal gyri among participants with childhood trauma (Lu et al., 2017). Decreased regional homogeneity in the right posterior temporal gyrus was also reported in a small study of adults (Philip, Kuras, et al., 2013). Tursich et al. (2015) studied 21 adults with PTSD related to childhood trauma, and observed an association between decreased rsFC of the left posterior temporal gyrus and more severe hyperarousal symptoms. In addition, gray matter volumetric changes in the superior temporal gyrus have been reported with PTSD (De Bellis et al., 2002) and MDD (Lorenzetti et al., 2009). Behaviorally, children who had experienced maltreatment were found to have delayed language skills, which are encoded in temporal gyri, compared to children without maltreatment experiences (Culp et al., 1991; McFadyen & Kitson, 1996; Sylvestre et al., 2016). Our findings may provide a biological accounting for these reported language deficits.

Cluster 2 also contained the right insula. The insular cortex subserves different functions, including sensorimotor processing, emotional processing, and cognitive functioning (Uddin, Nomi, Hebert-Seropian, Ghaziri, & Boucher, 2017). It is subdivided into the anterior insula and the posterior insula. Based on the coordinates in MNI152 space, the identified region was the right posterior insula. The posterior insula has been functionally associated with primary and secondary somatomotor cortices, including the SMA, pre-SMA, and most of precentral and postcentral gyri (Deen, Pitskel, & Pelphrey, 2011). Zhang et al. (2016) observed that relative to controls (n=20), patients with PTSD (n=20) who experienced serious vehicle accidents showed decreased rsFC between the right posterior insula and the postcentral gyrus. These reported results are consistent with our current finding that the right posterior insula and postcentral gyrus exhibited decreased functional connectivity as TE number increased. We speculate that traumatic experiences may have an impact on the regulation of how somatosensory information is processed in the posterior insula, which may partially account for the strong connection between PTSD and altered emotional processing after traumatic experiences (Brewin & Holmes, 2003; Shepherd & Wild, 2014).

Clusters 3 and 4 included sensorimotor regions, including the left SMA, paracentral lobule, precentral and postcentral gyri. The disruption of rsFC in these sensorimotor regions has not been reported in non-psychiatric individuals with traumatic experiences. For patients with PTSD, both structural and functional changes in sensorimotor regions have been reported. Sui et al. (2010) observed increased gray matter density in the postcentral cortex, and Zhang et al. (2016) observed decreased functional connectivity between the right posterior insula and the postcentral gyrus in patients with PTSD, compared to controls. For patients with MDD, decreased regional homogeneity has been observed in the left postcentral gyrus and left precentral gyrus compared to controls (Liu et al., 2010). As mentioned above, the posterior insula is functionally connected to these sensorimotor regions. Taken together, these results suggest that the processing of somatosensory information could be impacted by traumatic experiences. Furthermore, sensorimotor circuits subserve the comprehension of phonological information, semantic categories, and grammar (Pulvermüller & Fadiga, 2010), suggesting that rsFC changes in these sensorimotor regions could have an impact on language skills.

The resting-state networks defined by using the identified clusters as seeds highly overlapped with the somatomotor network. Behroozmand et al. (2015) identified a network involved in speech production and motor control in an fMRI study. This network included the bilateral superior temporal gyri, Heschl’s gyrus, precentral gyrus, SMA, Rolandic operculum, postcentral gyrus, and the right inferior frontal gyrus, which are concordant with the regions in Network 1, indicating that speech production and motor control might be affected by traumatic experiences in youth. These results suggest that traumatic experiences might have an impact on rsFC in youth, even in the absence of concurrent psychiatric disorders. Furthermore, the language deficits observed in children with maltreatment (Culp et al., 1991; McFadyen & Kitson, 1996; Sylvestre et al., 2016) could be explained by decreased rsFC in the somatomotor network.

### The interaction between sex and TE number

We identified four clusters that showed significant sex differences in the association between TE number and rsFC. These clusters were mainly located within the DMN. In addition, the resting-state networks defined by using these clusters as seeds also highly overlapped with the DMN. The DMN is a large intrinsic functional network. Its functions are associated with self-oriented processing (Gusnard, Akbudak, Shulman, & Raichle, 2001) and preparing for responses to environmental stimuli (Raichle & Gusnard, 2005). Numerous studies have reported decreased rsFC in the DMN among non-psychiatric individuals with trauma exposure (Lu et al., 2017; Philip, Sweet, et al., 2013) and in patients with PTSD relative to controls (Bluhm et al., 2009; Lanius et al., 2010; Tursich et al., 2015). However, sex-specific differences in rsFC have not been well-documented. Our analyses used a large youth population dataset with 57.8% females, providing power to identify significant sex-specific associations. We observed that within these identified clusters, females showed decreased functional connectivity, whereas males showed increased connectivity as TE number increased. Jointly considering the current results with the previous finding of decreased rsFC within the DMN in PTSD patients (Bluhm et al., 2009; Lanius et al., 2010; Tursich et al., 2015), we infer that females may be more susceptible to developing PTSD following traumatic exposure. This view is consistent with the finding that female victims are more likely to develop PTSD after experiencing trauma (Breslau, 2009; Tolin & Foa, 2008). The current findings offer a potential neural explanation for the higher risk in females and a potential biomarker for monitoring therapeutic trials.

## Limitations

There are several limitations with the current study. First, the traumatic experience assessment was based on questions in a self-reported PTSD scale, instead of a standard scale to measure childhood trauma, such as the Childhood Trauma Questionnaire (Bernstein et al., 2003). Self-reported traumatic experiences might not reflect actual traumatic exposure. More stringent measurements are needed in future studies. Second, information regarding traumatic experiences is limited. We only obtained the number of categories of TE, but did not evaluate the trauma type, timing, duration, or severity. Sex-specific associations observed in this study may have arisen due to different trauma types experienced by males and females. Further studies are needed to verify our results and disentangle the association between these specific aspects of traumatic experiences and rsFC. Third, although the PNC participants were not seeking help for psychiatric issues, some of the participants in our study might have developed PTSD or other mental disorders. These participants may have distorted our results, as we aimed to test our hypotheses in a non-psychiatric population. Finally, we cannot infer causality based on this cross-sectional dataset, so it is difficult to attribute differences in rsFC specifically to traumatic experiences. Longitudinal data are needed to analyze the trajectories of change in rsFC following traumatic experiences. It is also worth noting that it remains unclear whether the sex-specific differences we observed were due to different reactions of the brain in coping with trauma or different levels of vulnerability to PTSD between males and females.

## Conclusions

In a large sample of non-psychiatric youth, the current study is the first to report a significant association between traumatic experiences and reduced rsFC within the somatomotor network, as well as sex-specific associations between traumatic experiences and rsFC within the DMN. The disturbances in these brain networks provide a possible neural basis for the language deficits observed in children following traumatic exposures and the higher risk for females to develop PTSD compared to males. Further research is needed to establish the causality of these associations and to evaluate how these associations are qualified by trauma type, timing, duration, and severity.

## Key points

- Traumatic experiences are a significant risk factor for multiple psychiatric disorders and adversely affect cognitive functions. Little is known about the sex-specific effects of traumatic experiences on functional connectivity.
- In a large sample of non-psychiatric youth, this study identified an association between traumatic experiences and reduced resting-state functional connectivity within the somatomotor network.
- This study also identified sex-specific associations between traumatic experiences and resting-state functional connectivity within the default mode network.
- The current findings offer a possible neural basis for the language deficits observed in children following traumatic exposures and the higher risk for females to develop PTSD compared to males.

## Acknowledgements

This project was supported by the Manton Foundation (JGM and JRG), the Eunice Kennedy Shriver National Institute of Child Health and Human Development (P50HD027802 to JGM and JRG), the National Human Genome Research Institute (R01HG010171 to JRG), and U.S. National Institutes of Health (R01HG010171 and R01MH116527 to HZ). The content is solely the responsibility of the authors and does not necessarily represent the official views of the National Institutes of Health.

Support for the collection of the datasets was provided by grant RC2MH089983 awarded to Raquel Gur and RC2MH089924 awarded to Hakon Hakonarson. All participants were recruited through the Center for Applied Genomics at The Children’s Hospital in Philadelphia. The PNC datasets used in this manuscript were obtained from dbGaP at https://www.ncbi.nlm.nih.gov/projects/gap/cgi-bin/study.cgi?study_id=phs000607.v2.p2 through dbGaP accession <u>phs000607.v2.p2</u>.

## References

Association, A. P. (2013). Diagnostic and statistical manual of mental disorders (DSM-5®): American Psychiatric Pub.

Barzilay, R., Calkins, M. E., Moore, T. M., Wolf, D. H., Satterthwaite, T. D., Scott, J. C., … Gur, R. E. (2019). Association between traumatic stress load, psychopathology, and cognition in the Philadelphia Neurodevelopmental Cohort. Psychological medicine, 49(2), 325–334.

Behroozmand, R., Shebek, R., Hansen, D. R., Oya, H., Robin, D. A., Howard III, M. A., & Greenlee, J. D. (2015). Sensory–motor networks involved in speech production and motor control: An fMRI study. Neuroimage, 109, 418–428.

Bernstein, D. P., Stein, J. A., Newcomb, M. D., Walker, E., Pogge, D., Ahluvalia, T., … Desmond, D. (2003). Development and validation of a brief screening version of the Childhood Trauma Questionnaire. Child abuse & neglect, 27(2), 169–190.

Bluhm, R. L., Williamson, P. C., Osuch, E. A., Frewen, P. A., Stevens, T. K., Boksman, K., … Lanius, R. A. (2009). Alterations in default network connectivity in posttraumatic stress disorder related to early-life trauma. Journal of psychiatry & neuroscience: JPN, 34(3), 187.

Breslau, N. (2009). The epidemiology of trauma, PTSD, and other posttrauma disorders. Trauma, Violence, & Abuse, 10(3), 198–210.

Brewin, C. R., & Holmes, E. A. (2003). Psychological theories of posttraumatic stress disorder. Clinical psychology review, 23(3), 339–376.

Briere, J., & Elliott, D. M. (2003). Prevalence and psychological sequelae of self-reported childhood physical and sexual abuse in a general population sample of men and women. Child abuse & neglect, 27(10), 1205–1222.

Calkins, M. E., Moore, T. M., Merikangas, K. R., Burstein, M., Satterthwaite, T. D., Bilker, W. B., … Mentch, F. (2014). The psychosis spectrum in a young US community sample: findings from the Philadelphia Neurodevelopmental Cohort. World Psychiatry, 13(3), 296–305.

Copeland, W. E., Keeler, G., Angold, A., & Costello, E. J. (2007). Traumatic events and posttraumatic stress in childhood. Archives of general psychiatry, 64(5), 577–584.

Cox, R. W. (1996). AFNI: software for analysis and visualization of functional magnetic resonance neuroimages. Computers and Biomedical research, 29(3), 162–173.

Culp, R. E., Watkins, R. V., Lawrence, H., Letts, D., Kelly, D. J., & Rice, M. L. (1991). Maltreated children’s language and speech development: Abused, neglected, and abused and neglected. First Language, 11(33), 377–389.

De Bellis, M. D., Keshavan, M. S., Frustaci, K., Shifflett, H., Iyengar, S., Beers, S. R., & Hall, J. (2002). Superior temporal gyrus volumes in maltreated children and adolescents with PTSD. Biological psychiatry, 51(7), 544–552.

Deen, B., Pitskel, N. B., & Pelphrey, K. A. (2011). Three systems of insular functional connectivity identified with cluster analysis. Cerebral cortex, 21(7), 1498–1506.

Felmingham, K., Williams, L. M., Kemp, A. H., Liddell, B., Falconer, E., Peduto, A., & Bryant, R. (2010). Neural responses to masked fear faces: sex differences and trauma exposure in posttraumatic stress disorder. Journal of abnormal psychology, 119(1), 241.

Gusnard, D. A., Akbudak, E., Shulman, G. L., & Raichle, M. E. (2001). Medial prefrontal cortex and self-referential mental activity: relation to a default mode of brain function. Proceedings of the National Academy of Sciences, 98(7), 4259–4264.

Heim, C., & Nemeroff, C. B. (2001). The role of childhood trauma in the neurobiology of mood and anxiety disorders: preclinical and clinical studies. Biological psychiatry, 49(12), 1023–1039.

Kaufman, J., Birmaher, B., Brent, D., Rao, U., Flynn, C., Moreci, P., … Ryan, N. (1997). Schedule for affective disorders and schizophrenia for school-age children-present and lifetime version (K-SADS-PL): initial reliability and validity data. Journal of the American Academy of Child & Adolescent Psychiatry, 36(7), 980–988.

Kilpatrick, D. G., Ruggiero, K. J., Acierno, R., Saunders, B. E., Resnick, H. S., & Best, C. L. (2003). Violence and risk of PTSD, major depression, substance abuse/dependence, and comorbidity: results from the National Survey of Adolescents. Journal of consulting and clinical psychology, 71(4), 692.

Lanius, R., Bluhm, R., Coupland, N., Hegadoren, K., Rowe, B., Theberge, J., … Brimson, M. (2010). Default mode network connectivity as a predictor of post-traumatic stress disorder symptom severity in acutely traumatized subjects. Acta Psychiatrica Scandinavica, 121(1), 33–40.

Liu, Z., Xu, C., Xu, Y., Wang, Y., Zhao, B., Lv, Y., … Du, C. (2010). Decreased regional homogeneity in insula and cerebellum: a resting-state fMRI study in patients with major depression and subjects at high risk for major depression. Psychiatry Research: Neuroimaging, 182(3), 211–215.

Lorenzetti, V., Allen, N. B., Fornito, A., & Yücel, M. (2009). Structural brain abnormalities in major depressive disorder: a selective review of recent MRI studies. Journal of affective disorders, 117(1-2), 1–17.

Lu, S., Gao, W., Wei, Z., Wang, D., Hu, S., Huang, M., … Li, L. (2017). Intrinsic brain abnormalities in young healthy adults with childhood trauma: A resting-state functional magnetic resonance imaging study of regional homogeneity and functional connectivity. Australian & New Zealand Journal of Psychiatry, 51(6), 614–623.

Majer, M., Nater, U. M., Lin, J.-M. S., Capuron, L., & Reeves, W. C. (2010). Association of childhood trauma with cognitive function in healthy adults: a pilot study. BMC neurology, 10(1), 61.

Marusak, H. A., Etkin, A., & Thomason, M. E. (2015). Disrupted insula-based neural circuit organization and conflict interference in trauma-exposed youth. NeuroImage: Clinical, 8, 516–525.

McFadyen, R. G., & Kitson, W. J. (1996). Language comprehension and expression among adolescents who have experienced childhood physical abuse. Journal of Child Psychology and Psychiatry, 37(5), 551–562.

Pérez-Fuentes, G., Olfson, M., Villegas, L., Morcillo, C., Wang, S., & Blanco, C. (2013). Prevalence and correlates of child sexual abuse: a national study. Comprehensive psychiatry, 54(1), 16–27.

Philip, N. S., Kuras, Y. I., Valentine, T. R., Sweet, L. H., Tyrka, A. R., Price, L. H., & Carpenter, L. L. (2013). Regional homogeneity and resting state functional connectivity: associations with exposure to early life stress. Psychiatry Research: Neuroimaging, 214(3), 247–253.

Philip, N. S., Sweet, L. H., Tyrka, A. R., Price, L. H., Bloom, R. F., & Carpenter, L. L. (2013). Decreased default network connectivity is associated with early life stress in medication-free healthy adults. European Neuropsychopharmacology, 23(1), 24–32.

Pulvermüller, F., & Fadiga, L. (2010). Active perception: sensorimotor circuits as a cortical basis for language. Nature reviews neuroscience, 11(5), 351–360.

Raichle, M. E., & Gusnard, D. A. (2005). Intrinsic brain activity sets the stage for expression of motivated behavior. Journal of Comparative Neurology, 493(1), 167–176.

Saad, Z. S., Reynolds, R. C., Argall, B., Japee, S., & Cox, R. W. (2004). SUMA: an interface for surface-based intra-and inter-subject analysis with AFNI. Paper presented at the 2004 2nd IEEE International Symposium on Biomedical Imaging: Nano to Macro (IEEE Cat No. 04EX821).

Satterthwaite, T. D., Connolly, J. J., Ruparel, K., Calkins, M. E., Jackson, C., Elliott, M. A., … Behr, M. (2016). The Philadelphia Neurodevelopmental Cohort: A publicly available resource for the study of normal and abnormal brain development in youth. Neuroimage, 124, 1115–1119.

Satterthwaite, T. D., Elliott, M. A., Ruparel, K., Loughead, J., Prabhakaran, K., Calkins, M. E., … Riley, M. (2014). Neuroimaging of the Philadelphia neurodevelopmental cohort. Neuroimage, 86, 544–553.

Scheinost, D., Benjamin, J., Lacadie, C., Vohr, B., Schneider, K. C., Ment, L. R., … Constable, R. T. (2012). The intrinsic connectivity distribution: a novel contrast measure reflecting voxel level functional connectivity. Neuroimage, 62(3), 1510–1519.

Shepherd, L., & Wild, J. (2014). Emotion regulation, physiological arousal and PTSD symptoms in trauma-exposed individuals. Journal of behavior therapy and experimental psychiatry, 45(3), 360–367.

Shvil, E., Sullivan, G. M., Schafer, S., Markowitz, J. C., Campeas, M., Wager, T. D., … Neria, Y. (2014). Sex differences in extinction recall in posttraumatic stress disorder: a pilot fMRI study. Neurobiology of learning and memory, 113, 101–108.

Sripada, R. K., King, A. P., Welsh, R. C., Garfinkel, S. N., Wang, X., Sripada, C. S., & Liberzon, I. (2012). Neural dysregulation in posttraumatic stress disorder: evidence for disrupted equilibrium between salience and default mode brain networks. Psychosomatic medicine, 74(9), 904.

Sui, S. G., Wu, M. X., King, M. E., Zhang, Y., Ling, L., Xu, J. M., … Li, L. J. (2010). Abnormal grey matter in victims of rape with PTSD in Mainland China: a voxel-based morphometry study. Acta neuropsychiatrica, 22(3), 118–126.

Sylvestre, A., Bussières, È.-L., & Bouchard, C. (2016). Language problems among abused and neglected children: A meta-analytic review. Child maltreatment, 21(1), 47–58.

Thomas Yeo, B., Krienen, F. M., Sepulcre, J., Sabuncu, M. R., Lashkari, D., Hollinshead, M., … Polimeni, J. R. (2011). The organization of the human cerebral cortex estimated by intrinsic functional connectivity. Journal of neurophysiology, 106(3), 1125–1165.

Tolin, D. F., & Foa, E. B. (2008). Sex differences in trauma and posttraumatic stress disorder: a quantitative review of 25 years of research.

Tursich, M., Ros, T., Frewen, P., Kluetsch, R., Calhoun, V. D., & Lanius, R. A. (2015). Distinct intrinsic network connectivity patterns of post-traumatic stress disorder symptom clusters. Acta Psychiatrica Scandinavica, 132(1), 29–38.

Uddin, L. Q., Nomi, J. S., Hebert-Seropian, B., Ghaziri, J., & Boucher, O. (2017). Structure and function of the human insula. Journal of clinical neurophysiology: official publication of the American Electroencephalographic Society, 34(4), 300.

Viard, A., Mutlu, J., Chanraud, S., Guenolé, F., Egler, P.-J., Gérardin, P., … Guillery-Girard, B. (2019). Altered default mode network connectivity in adolescents with post-traumatic stress disorder. NeuroImage: Clinical, 22, 101731.

Zhang, Y., Xie, B., Chen, H., Li, M., Guo, X., & Chen, H. (2016). Disrupted resting-state insular subregions functional connectivity in post-traumatic stress disorder. Medicine, 95(27).

