## Supplementary information for "Sex-specific Associations Between Traumatic Experiences and Resting-state Functional Connectivity in the Philadelphia Neurodevelopmental Cohort"

**Appendix S1. Traumatic experience assessment**

Nine questions from the PTSD assessment scale (Table S1) in GOASSESS (Calkins et al., 2014) were used to assess traumatic experiences. Responses for each question were assigned a value of 1 for yes, and 0 for no. Questions 4 and 5 probed sexual assault. Most subjects left question 5 unanswered, which asked about rape, so we combined questions 4 and 5 to capture as much information as possible about sexual assault. We assigned a value of 1 when the answer to either question 4 or 5 was yes. Then, for each subject, we summed the number of unweighted traumatic events for use in subsequent regression analyses.

**Appendix S2. Image acquisition**

MRI images were collected using a single 3T Siemens TIM Trio whole-body scanner located in the Hospital of the University of Pennsylvania (Satterthwaite et al., 2014). Anatomical images were acquired using a magnetization prepared, rapid-acquisition gradient-echo (MPRAGE) sequence (matrix size=256x192; voxel size=0.9x0.9x1 mm; field-of-view (FoV)=180x240 mm; repetition time (TR)=1810 ms; echo time (TE)=3.5 ms; flip angle=9$^{\circ}$). Resting-state BOLD scans were obtained using a single-shot, interleaved multi-slice, gradient-echo, echo planar imaging (GE-EPI) sequence (matrix size=64x64; voxel size=3x3x3 mm; FoV=192x192 mm; TR=3000 ms; TE=32 ms; flip angle=90$^{\circ}$). The total duration of the resting-state scan was 6.2 min. Subjects were asked to keep their eyes open, stay awake, fixate on the displayed crosshair, and remain still.

**Appendix S3. Tissue-based regression**

Tissue-based regression was performed to regress out the average signals of individual eroded white matter masks and the first three principal components of individual eroded lateral ventricle masks using the “fast” ANATICOR program in AFNI (Jo, Saad, Simmons, Milbury, & Cox, 2010). Freesurfer (Dale, Fischl, & Sereno, 1999) was used to generate white matter and lateral ventricle masks for each participant based on anatomical scans and using the Desikan-Killiany atlas (Desikan et al., 2006).

**Appendix S4. Intrinsic connectivity distribution (ICD)**

ICD is a method to measure voxel-level functional connectivity as described by Scheinost et al. (2012). Correlation coefficients were calculated between each voxel and every other voxel in the brain using time course residuals from the general linear model. Then, positive correlation coefficients were used to generate a histogram. The survival function computed from this histogram was approximated using a Weibull distribution model $S\left( r \right)=e^{-\alpha*r^{\beta}}$. The survival function was characterized by a variance parameter alpha and a shape parameter beta. Smaller alpha values and larger beta values for any voxel represent a higher density of strong connections with other voxels.

**Table S1. Questions to assess traumatic experiences**

|  | **Questions** |
| --- | --- |
| 1 | Have you ever been in a flood or a tornado or an earthquake or a hurricane or some other natural disaster where you thought you were going to die or be seriously hurt? |
| 2 | Have you ever been in a situation where you thought you or someone close to you was going to be killed or be hurt very badly? |
| 3 | Have you ever been attacked by somebody or badly beaten? |
| 4 | Have you ever been very upset by someone forcing you to do something sexual? |
| 5 | Have you ever been attacked sexually or raped? |
| 6 | Have you ever been threatened with a weapon? |
| 7 | Have you ever been in a bad accident? |
| 8 | Other than television or at the movies, have you ever seen or heard somebody get killed or get hurt very badly or die? |
| 9 | Have you ever been very upset by seeing a dead body or by seeing pictures of the dead body of somebody you knew well? |

**Table S2. Demographic information for participants with and without traumatic exposure**

| **Characteristics** | **Non-exposure** **(n=477)** | **Exposure** **(n=437)** | **Statistics (ANOVA** $\boldsymbol{F}$**/ Pearson** $\mathbf{X}^{\mathbf{2}}$**)** | ***P* value** |
| --- | --- | --- | --- | --- |
| Female sex, No. (%) | 274 (57.4) | 255 (58.4) | $X^{2}$=0.045 | 0.83 |
| Race |  |  | $X^{2}$=19.7 | <0.001 |
| African-American, No. (%) | 187 (39.2) | 236 (54.0) |  |  |
| European-American, No. (%) | 233 (48.8) | 163 (37.3) |  |  |
| Other, No. (%) | 54 (11.3) | 37 (8.5) |  |  |
| Age, mean (SD), y | 13.9 (3.50) | 15.3 (2.94) | *F*=39.7 | <0.001 |
| ME, mean (SD), y | 14.4 (2.41) | 13.9 (2.42) | *F*=10.4 | 0.001 |

Abbreviation: ME, years of maternal education

**Table S3. Exposure of participants to each category of traumatic event**

| **Traumatic event** | **N (% of cohort)** | **Number of females in category (%)** |
| --- | --- | --- |
| 1. Experienced a natural disaster | 31 (3.4) | 20 (64.5) |
| 2. Thought you or someone close to you was going to be killed or be hurt very badly | 120 (13.1) | 66 (55.0) |
| 3. Attacked by somebody or badly beaten | 55 (6.0) | 26 (47.3) |
| 4. Sexually forced | 26 (2.8) | 22 (84.6) |
| 5. Threatened with a weapon | 60 (6.6) | 17 (28.4) |
| 6. Experienced a bad accident | 91 (10.0) | 52 (57.1) |
| 7. Witnessed someone getting killed, badly beaten, or die | 211 (23.1) | 124 (58.8) |
| 8. Upset by seeing a dead body or pictures of the dead body of somebody you knew well | 228 (24.9) | 137 (60.1) |

**Table S4. Brain regions in resting-state Network 2**

| **Hemisphere** | **Regions** |
| --- | --- |
| Bilateral | superior temporal gyrus, Heschl’s gyrus, SMA, middle cingulate cortex, precentral gyrus, postcentral gyrus, Rolandic operculum, insula |
| Right | paracentral lobule |

**Table S5. Brain regions in resting-state Network 3**

| **Hemisphere** | **Regions** |
| --- | --- |
| Bilateral | SMA, paracentral lobule, precentral gyrus, postcentral gyrus |

**Table S6. Brain regions in resting-state Network 4**

| **Hemisphere** | **Regions** |
| --- | --- |
| Bilateral | paracentral lobule, precentral gyrus, postcentral gyrus, SMA |

**Table S7. Brain regions in resting-state Network 6**

| **Hemisphere** | **Regions** |
| --- | --- |
| Bilateral | precuneus, middle cingulate cortex, cuneus, posterior cingulate cortex, angular gyrus |
| Left | middle occipital gyrus |
| Right | superior frontal gyrus, middle frontal gyrus, mid-orbital gyrus, anterior cingulate cortex |

**Table S8. Brain regions in resting-state Network 7**

| **Hemisphere** | **Regions** |
| --- | --- |
| Bilateral | precuneus, middle cingulate cortex, posterior cingulate cortex, mid-orbital gyrus, angular gyrus, anterior cingulate cortex |

**Table S9. Brain regions in resting-state Network 8**

| **Hemisphere** | **Regions** |
| --- | --- |
| Bilateral | precuneus, middle cingulate cortex, posterior cingulate cortex, anterior cingulate cortex, mid-orbital gyrus, angular gyrus |
| Left | superior frontal gyrus, middle frontal gyrus, middle occipital gyrus |

**Fig S1. Peak nodes within Network 1**

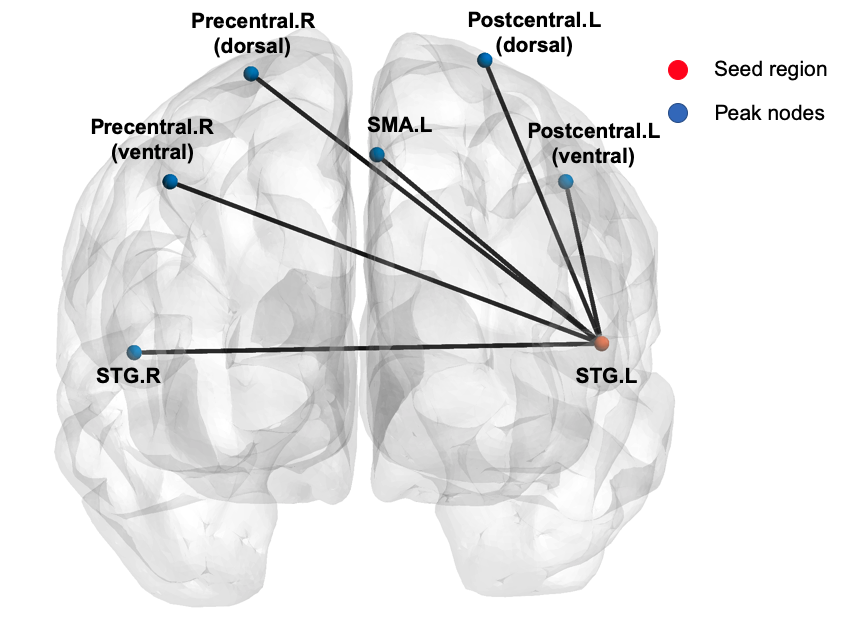

Peak nodes whose resting-state time courses were highly correlated with Cluster 1 within Network 1. Peak nodes were extracted using AFNI (Cox, 1996) (*3dExtrema*; minimum separation distance of 30 mm, or 10 voxels). BrainNet Viewer (Xia, Wang, & He, 2013) was used for visualization. STG: superior temporal gyrus, SMA: supplementary motor area, R: right hemisphere, L: left hemisphere.

**Fig S2. Resting-state Network 2**

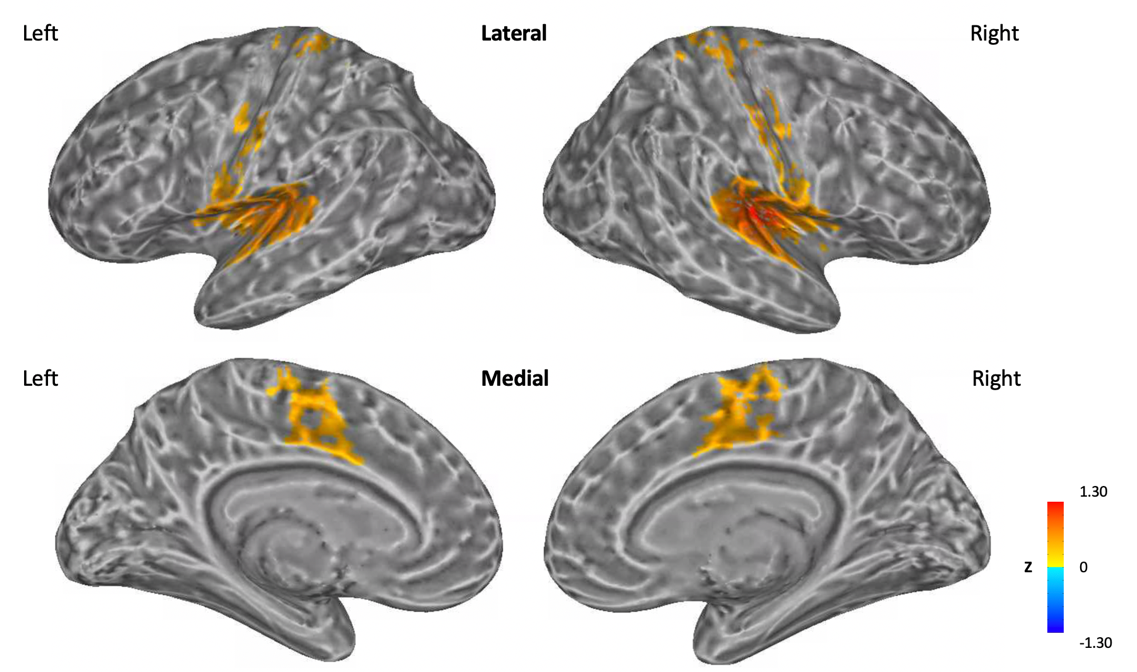

Resting-state Network 2 defined by using Cluster 2 as a seed region. The correlation coefficients between the average time-series data of the seed region with every other voxel were calculated and transformed to z scores. A *t*-test identified significant regions (p < 1x10^-44^, FDR < 3x10^-16^) that defined a resting-state network. Colors represent the value of z scores within the defined resting-state network.

**Fig S3. Resting-state Network 3**

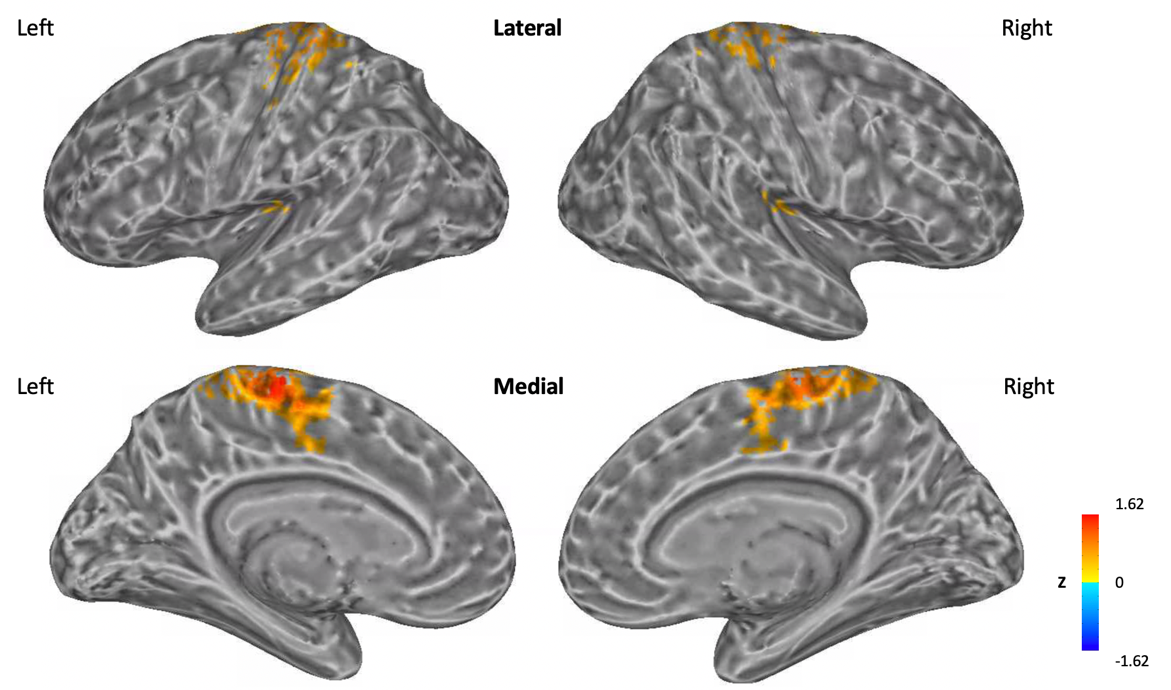

Resting-state Network 3 defined by using Cluster 3 as a seed region. The correlation coefficients between the average time-series data of the seed region with every other voxel were calculated and transformed to z scores. A *t*-test identified significant regions (p < 1x10^-44^, FDR < 3x10^-16^) that defined a resting-state network. Colors represent the value of z scores within the defined resting-state network.

**Fig S4. Resting-state Network 4**

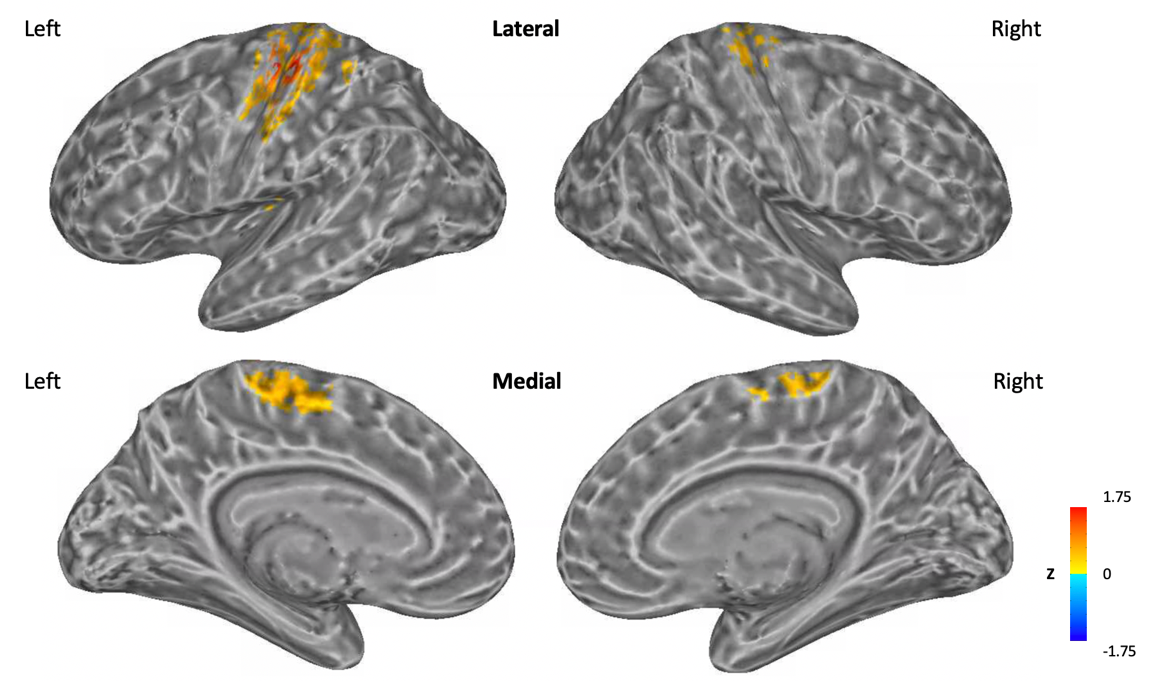

Resting-state Network 4 defined by using Cluster 4 as a seed region. The correlation coefficients between the average time-series data of the seed region with every other voxel were calculated and transformed to z scores. A *t*-test identified significant regions (p < 1x10^-44^, FDR < 3x10^-16^) that defined a resting-state network. Colors represent the value of z scores within the defined resting-state network.

**Fig S5. Peak nodes within Network 5**

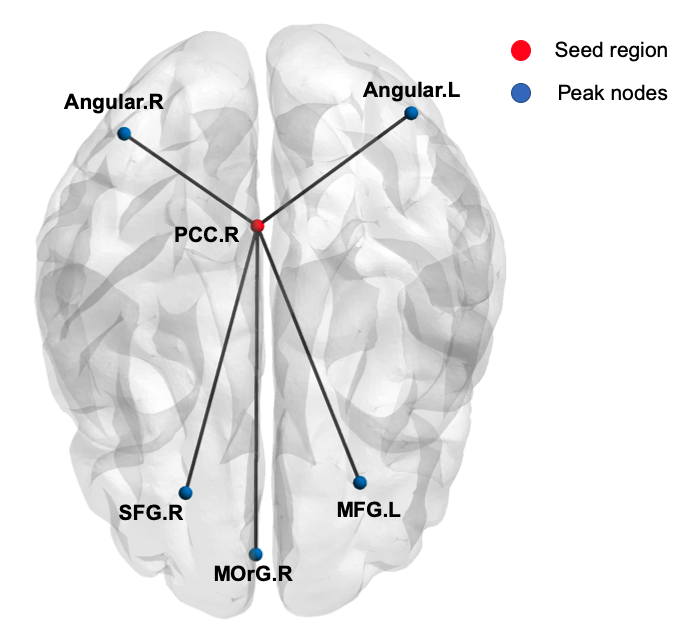

Peak nodes whose resting-state time courses were highly correlated with Cluster 5 within Network 5. Peak nodes were extracted using AFNI (Cox, 1996) (*3dExtrema*; minimum separation distance of 30 mm, or 10 voxels). BrainNet Viewer (Xia et al., 2013) was used for visualization. PCC: posterior cingulate cortex, MFG: middle frontal gyrus, SFG: superior frontal gyrus, MOrG: medial-orbital gyrus, R: right hemisphere, L: left hemisphere.

**Fig S6. Resting-state Network 6**

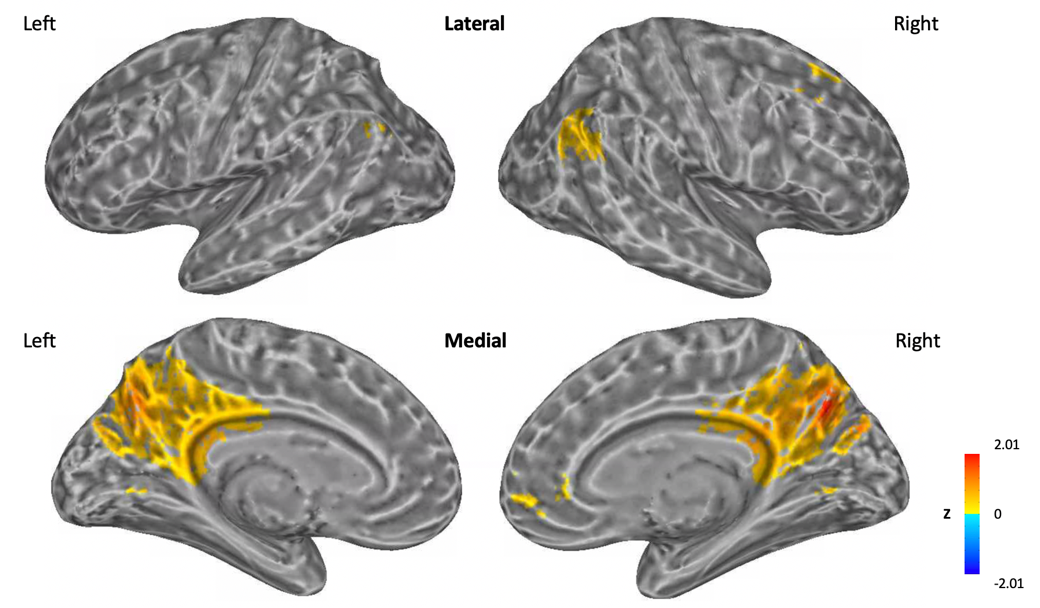

Resting-state Network 6 defined by using Cluster 6 as a seed region. The correlation coefficients between the average time-series data of the seed region with every other voxel were calculated and transformed to z scores. A *t*-test identified significant regions (p < 1x10^-44^, FDR < 3x10^-16^) that defined a resting-state network. Colors represent the value of z scores within the defined resting-state network.

**Fig S7. Resting-state Network 7**

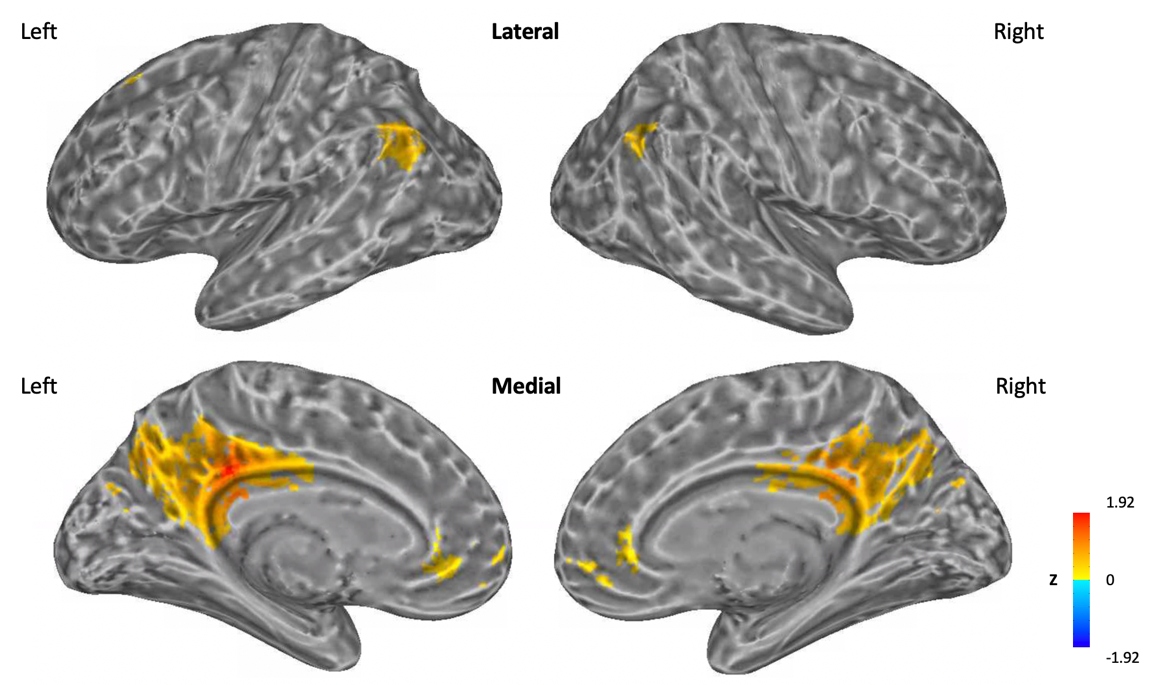

Resting-state Network 7 defined by using Cluster 7 as a seed region. The correlation coefficients between the average time-series data of the seed region with every other voxel were calculated and transformed to z scores. A *t*-test identified significant regions (p < 1x10^-44^, FDR < 3x10^-16^) that defined a resting-state network. Colors represent the value of z scores within the defined resting-state network.

**Fig S8. Resting-state Network 8**

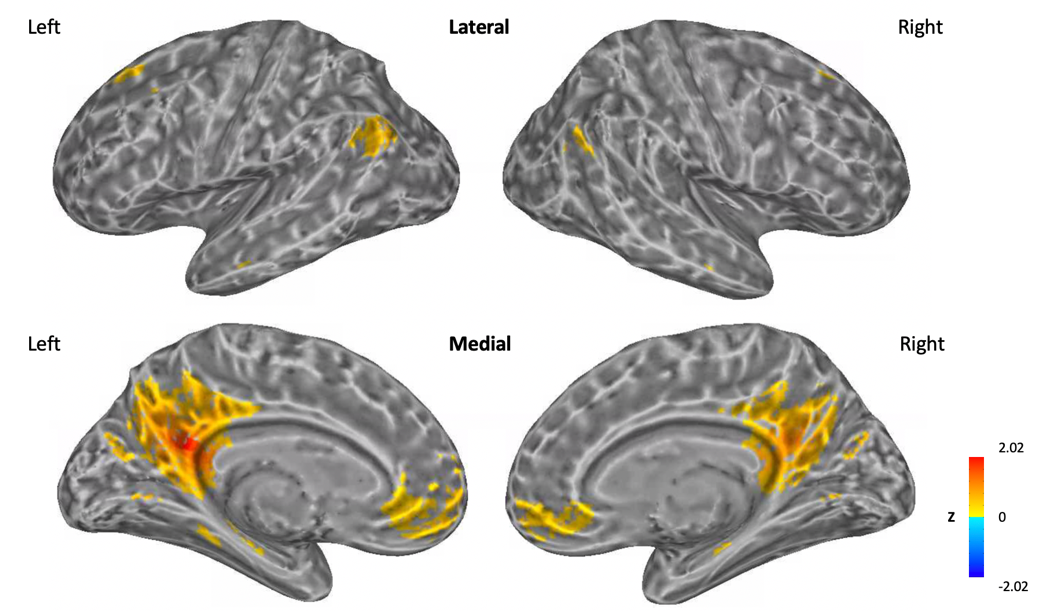

Resting-state Network 8 defined by using Cluster 8 as a seed region. The correlation coefficients between the average time-series data of the seed region with every other voxel were calculated and transformed to z scores. A *t*-test identified significant regions (p < 1x10^-44^, FDR < 3x10^-16^) that defined a resting-state network. Colors represent the value of z scores within the defined resting-state network.

**References**

Calkins, M. E., Moore, T. M., Merikangas, K. R., Burstein, M., Satterthwaite, T. D., Bilker, W. B., . . . Mentch, F. (2014). The psychosis spectrum in a young US community sample: findings from the Philadelphia Neurodevelopmental Cohort. *World Psychiatry, 13*(3), 296-305.

Cox, R. W. (1996). AFNI: software for analysis and visualization of functional magnetic resonance neuroimages. *Computers and Biomedical research, 29*(3), 162-173.

Dale, A. M., Fischl, B., & Sereno, M. I. (1999). Cortical surface-based analysis: I. Segmentation and surface reconstruction. *Neuroimage, 9*(2), 179-194.

Desikan, R. S., Ségonne, F., Fischl, B., Quinn, B. T., Dickerson, B. C., Blacker, D., . . . Hyman, B. T. (2006). An automated labeling system for subdividing the human cerebral cortex on MRI scans into gyral based regions of interest. *Neuroimage, 31*(3), 968-980.

Jo, H. J., Saad, Z. S., Simmons, W. K., Milbury, L. A., & Cox, R. W. (2010). Mapping sources of correlation in resting state FMRI, with artifact detection and removal. *Neuroimage, 52*(2), 571-582.

Satterthwaite, T. D., Elliott, M. A., Ruparel, K., Loughead, J., Prabhakaran, K., Calkins, M. E., . . . Riley, M. (2014). Neuroimaging of the Philadelphia neurodevelopmental cohort. *Neuroimage, 86*, 544-553.

Scheinost, D., Benjamin, J., Lacadie, C., Vohr, B., Schneider, K. C., Ment, L. R., . . . Constable, R. T. (2012). The intrinsic connectivity distribution: a novel contrast measure reflecting voxel level functional connectivity. *Neuroimage, 62*(3), 1510-1519.

Xia, M., Wang, J., & He, Y. (2013). BrainNet Viewer: a network visualization tool for human brain connectomics. *PloS one, 8*(7), e68910.
